# ER-phagy and mitophagy programs in human CD4+ memory T cells before and after activation

**DOI:** 10.1101/2025.11.25.690386

**Authors:** Diego Morone, Andrea Raimondi, Maurizio Molinari

**Affiliations:** Università della Svizzera italiana (USI), Faculty of Biomedical Sciences, Institute for Research in Biomedicine, CH-6500 Bellinzona, Switzerland; Graduate School for Cellular and Biomedical Sciences, University of Bern, CH-3000, Bern, Switzerland; School of Life Sciences, École Polytechnique Fédérale de Lausanne, CH-1015 Lausanne, Switzerland

**Keywords:** CD4+ T cells, Endoplasmic reticulum (ER), ER-phagy, Endolysosomes (ELs), ER-phagy, Mitochondria, Mitophagy, Outer Mitochondrial Membrane (OMM)-phagy, Selective autophagy, T lymphocytes

## Abstract

T lymphocytes can remain in a resting state for very long times. Encountering their cognate antigen triggers their activation, leading to an increase in cell size, proliferation, enhanced protein production, and secretion of signaling molecules. This activation and clonal expansion phase is followed by a contraction phase, with a reduction in cell size, eventually leading to cell death or progression to resting (memory) cells. Here, we show that human resting primary CD4+ memory T cells degrade endoplasmic reticulum (ER) and mitochondrial portions via ER-phagy and mitophagy, respectively. The temporary interruption of ER and mitochondria turnover coupled with a transient induction of the unfolded protein response results in the enlargement of the ER and cellular volume. Cell growth is stabilized and then reversed during the clonal expansion and contraction phases by the resumption of a catabolic phase characterized by the reactivation of ER-phagy and outer mitochondrial membrane (OMM)-phagy. Collectively, our data show that the maintenance and the activity of human CD4+ memory T cells rely on finely tuned execution of anabolic and catabolic programs that regulate mass and activity of the ER and mitochondria.

## Introduction

T lymphocytes are fundamental components of adaptive immunity, the secondary line of defense against pathogens. T cells may remain in a resting state for a long time (1,2). Cells in this state are characterized by minimal net growth and basal nutrient uptake. Encountering their cognate antigen triggers their activation, leading to an increase in cell size, proliferation, enhanced protein production and secretion of signaling molecules (3). This is followed by a contraction phase, where cells stop to proliferate and reduce their size. Most of them eventually die, while some regain a resting state. Cells that have previously encountered their cognate antigen are called memory cells (1). Thymic selection generates two main populations of T cells, expressing CD4 or CD8 molecules. CD4+ T cells are key players in adaptive immune response, using specific effector function and orchestrating humoral responses to combat intracellular or extracellular pathogens (4).

The endoplasmic reticulum (ER) is the site of synthesis and maturation of proteins, lipids, and oligosaccharides (5). ER size and function are enhanced upon activation of unfolded protein response programs (UPRs) (6,7). The involvement of UPRs to enhance protein and lipid biosynthesis during cell differentiation (e.g., T cell or B cell activation) is well established in mice (8,9), plasma cells (10) and in human regulatory T cells (11,12). The energy required for this enhancement is obtained upon potentiation of the mass and activity of mitochondria, the organelles producing adenosine triphosphate (ATP) and plays an essential role in cellular metabolism and energy production (13). Thus, the *anabolic* programs enhancing ER and mitochondria mass and functions in T cells are well studied (8–12,14–16), as well as the consequences of their impaired activations (13,16–20).

Available studies also reveal the importance of *catabolic* programs in T cells biology, however, much less is known about the relevance of the controlled and selective lysosomal turnover of damaged, dysfunctional or excess ER and mitochondria in T cell activation and maintenance, as well as in the contraction phase that ensures returning of a fraction of the activated T cells to a resting, memory state.

ER-phagy is the selective autophagy of the ER. It is a conserved group of processes that control size and function of the ER, eliminate ER subdomains containing misfolded proteins and restore ER homeostasis after ER stress (21). It is induced to remodel the ER upon organogenesis and activation of cell differentiation programs (22–24). Notably, ER-phagy receptors, the membrane-associated proteins that control ER-phagy activation, are upregulated upon activation of CD8 T cells (25). Mitophagy is the selective autophagy of mitochondria. It relies on a series of selective autophagic processes that help maintain the mitochondrial fitness and/or to reduce the mitochondrial mass (26,27). ER-phagy and mitophagy are crucial for cells to adapt to external or internal stimuli, as they ensure the removal and replacement of damaged organelles with functional ones, thus maintaining cellular homeostasis. Defective autophagy, including defective ER-phagy and mitophagy, have been associated with persistence of defective mitochondria in CD4+ T cells as a result of defective mitochondrial turnover (28), thymocyte β-selection (29), CD8+ T cell exhaustion (30,31) and memory formation (32), T cell ageing (33), and myasthenia gravis (34). Recently, autophagy has been shown to be transiently suppressed in mouse CD8+ T cells after activation and *Listeria monocytogenes* infection *in vivo*, and to be repressed by cytokines during T cell differentiation (35). However, the temporal involvement of selective autophagy programs in human CD4+ memory T cells, particularly in relation to their resting and active states, remain poorly understood. Understanding the dynamics of selective autophagy pathways in human CD4+ T cells would provide additional insights on the complexity of T cell maintenance and regulation during activation.

Here, we monitor by imaging approaches, including imaging flow cytometry (IFC), laser scanning confocal microscopy (LSCM), and serial section Room Temperature-Transmission Electron Microscopy (ssRT-TEM), the fragmentation and lysosomal delivery of ER (ER-phagy), mitochondria (mitophagy), and OMM (OMM-phagy) in resting human CD4+ memory T cells, and in the same cells upon activation.

## Results

### Human memory CD4+ T cells transiently activate the UPR and enlarge the ER and the cytoplasm upon activation

T cells remain in a quiescent, resting state until encounter with cognate antigen. Upon activation, they start a clonal expansion phase, characterized by a high proliferation rate (among the highest in the cells of the human body (36)) and the production of cytokines driving their function (37). Flow cytometry and single-cell mechanical studies characterized the changes in cell size after T cell activation (38–40), and provided the confidence of large sample size and high-detail dynamics in single cell.

To extend these studies and offer a high-throughput microscopy characterization that combines a large sample size with measurement of the changes in cell and nucleus dimensions, we first assessed the morphological changes in cell and nuclear size upon T cell activation using IFC. To this end, CD4+ memory T cells were sorted and fixed before (**Fig. 1A**, upper row, Resting), or 1 to 5 days after incubation with CD3/CD28 antibody-coated plates and IL2-supplemented cell medium to mimic TCR clustering and costimulation (**Fig. 1A**, upper row, Day 1-Day 5). Sorting of total memory CD4+ T cells (i.e., central, effector, and terminally differentiated memory T cells) was performed from CD4-enriched peripheral blood mononuclear cells (PBMCs) of healthy donors with surface-directed antibodies as previously described (40). Cell were fixed with 4% paraformaldehyde (PFA) at days 1 to 5 post activation (p.a.). To determine cell and nuclear size, we employed IFC, which combines flow cytometry fluidics with a widefield microscopy setup and 2D image deconvolution, and enables rapid, high-throughput microscopic evaluation of subcellular parameters across a large sample size. Transmitted light and nuclear DNA staining (**Fig. 1A**, upper panels) showed a transient increase after activation (clonal expansion phase) of 2-fold in cells’ largest area from 79.0 ± 0.7 µm^2^ for resting cells, to 152.6 ± 1.8 µm^2^ for cells at day 2 p.a. (**Fig. 1B**). The experiment also revealed a transient 2-fold increase of the nuclei’s largest area from 34.5 ± 0.4 µm^2^ for resting cells, to 69.0 ± 1.2 µm^2^ at day 2 p.a. (**Fig. 1C**). Both the cells and nuclei area stopped expansion between days 2 and 3 p.a., to eventually start a contraction phase (**Figs. 1B-C**). Five days p.a., cells and nuclear size were still significantly bigger than before activation, with average sizes of 140.3 ± 1.2 and 60.7 ± 0.8 µm^2^, respectively (**Figs. 1B-C**).

**Figure 1.**
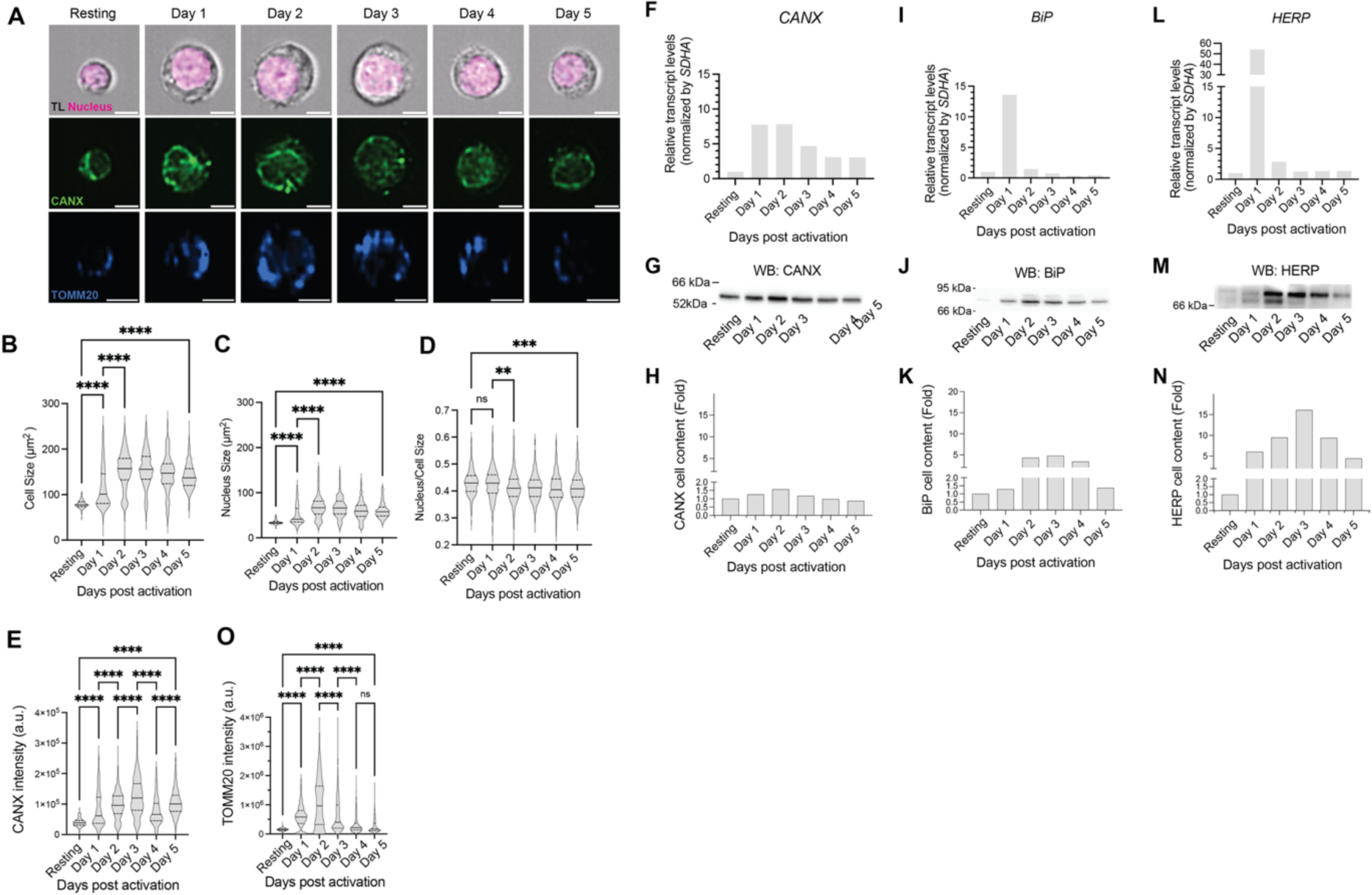
Human CD4+ memory T cells transiently increase ER and lysosomal content after activation. **A**) Representative images of CD4+ memory T cell during activation, as seen with transmitted light (TL) and stained with nuclear DNA staining, with ER membrane marker Calnexin (CANX) or with mitochondrial outer membrane marker TOMM20. Scale bars 7µm. **B**) Cell size, **C**) nucleus size, **D**) nucleus/cell size ratio (n>100 cells from 2 separate experiments). One-way ANOVA and Dunnett’s multiple comparisons test, ^ns^P>0.05, **P<0.01, ****P<0.0001. **E**) CANX intensity during activation (n>100 cells from 2 separate experiments). One-way ANOVA and Dunnett’s multiple comparisons test, ^ns^P>0.05, ****P<0.0001. **F**) RT-PCR of *CANX*, normalized by *SDHA*. **G**) Western Blot for total CANX content, normalized by number of cells. **H**) Quantification of a representative WB for CANX. **I**) RT-PCR of *BiP* transcripts, normalized by *SDHA*. **J**) Western Blot for total BiP content, normalized by number of cells. **K**) Quantification of a representative WB for BiP. **L**) RT-PCR of *HERP*, normalized by *SDHA*. **M**) Western Blot for total HERP content, normalized by number of cells. **N**) Quantification of a representative WB for HERP. **O**) TOMM20 intensity during activation (n>100 cells from 2 separate experiments). One-way ANOVA and Dunnett’s multiple comparisons test, ^ns^P>0.05, ****P<0.0001.

The activation of T cells relies on induction of an UPR that significantly enlarges the ER (8), which is site of proteins, lipids, and oligosaccharides synthesis. The enlargement of the ER, which extends the size of the cytoplasm thus increasing the ratio between cell and nuclear size (**Fig. 1D**), was confirmed by monitoring the variation in the intensity of the immunofluorescence associated with the ER-resident protein Calnexin (CANX, **Fig. 1A**, middle panel, **Fig. 1E**). The IFC revealed a transient increase in total CANX intensity up to 3-fold at day 3 p.a., which was followed by a decrease during the contraction phase to CANX levels that remained significantly higher than the CANX levels measured in resting cells at day 5 p.a. (**Fig. 1E**).

To verify if the enlargement of the ER mass observed by IFC was to ascribe to the induction of an UPR in our experimental set up, we examined the variations in transcript levels and total protein content of CANX (41), and of two other ER-resident reporters of ER stress induction, namely BiP (42,43) and HERP (44). Variations in CANX, BiP and HERP transcripts were measured with RT-PCR and normalized to *Succinate dehydrogenase complex subunit A* (*SDHA*), which is not transcriptionally induced upon CD4+ T cell activation (45). Variations in CANX, BiP and HERP protein levels were measured by Western Blot of detergent lysates of 1 · 10^6^ T cells collected before activation and at days 1-5 p.a..

The activation of T cells transiently increased the levels of CANX transcripts (7-fold at days 1 and 2 p.a., **Fig. 1F**) and protein (2-fold at day 2 p.a., **Figs. 1G-H**), consistent with the IFC data (**Fig. 1A**, middle row, and **Fig. 1E**). As expected upon induction of an UPR, also the transcript and protein levels of BiP (**Figs. 1I-1K**) and of HERP (**Figs. 1L-1N**) transiently increased, with higher maximal levels, but similar kinetics.

### Human memory CD4+ T cells transiently enhance the mitochondrial mass upon activation

T cell activation in presence of IL-2 induces mitochondrial fragmentation and increase in mitochondrial mass (14). To verify this, we measured, with IFC, the variation in the mitochondrial mass (i.e., in the intensity of the immunofluorescence associated with the OMM marker TOMM20) in the resting CD4+ cells and in the same cells upon activation (**Fig. 1A**, lower row). The values transiently increased by 8-fold, with a peak at day 2 p.a. (**Fig. 1O**) and a fast decrease that, in contrast to the ER situation, resulted in resumption of the levels measured in resting cells at days 4-5 p.a.

All in all, the increase in ER and mitochondrial content upon T cell activation, followed by a reduction in these organelles’ mass during the contraction phase, led us to verify the possible activation of catabolic phase during which ER and mitochondria are delivered to acidic degradative compartments for clearance by selective ER-phagy and mitophagy programs.

### Lysosomal delivery of ER portions in resting and in activated CD4+ memory T cells

To understand if the variations in ER and mitochondrial content observed in CD4+ memory T cells relies on activation of autophagic programs, we first examined ER delivery within LAMP1-positive degradative ELs in resting cells. To do so, resting T cells were treated for 12 hours with Bafilomycin A1 (BafA1) to inhibit the vacuolar-type proton-pump ATPase. Cell’s exposure to BafA1 neutralizes the lysosomal pH inactivates the acidic lysosomal hydrolases and results in the accumulation of undegraded material in the lumen of the degradative compartments (46). To perform a quantitative analysis of the accumulation of undegraded ER within the inactive T cells’ ELs, we developed a protocol for cytospinning and immunofluorescent staining of non-adherent cells. Briefly, cells were fixed and cytospinned on a round coverslip coated with Alcian blue, which was attached to the cytospin slide by water capillarity. After cytospin, the coverslip was detached with a few drops of water. Permeabilization and staining were performed as previously described (47). To quantify the presence of CANX-positive ER portions within LAMP1-positive EL, we adapted LysoQuant, a deep-learning quantitative approach that we developed for analyses of adherent cultured cells (48), for the automated quantification of lysosomal occupancy in non-adherent T cells. This adaptation consisted in a modification of the plugin pipeline to analyze T cells’ LAMP1-positive ELs, which are smaller than those found in immortalized cell lines, and to extend the pipeline to analyze 3D stacks.

Quantitative analyses of LCSM images revealed the accumulation of CANX-positive ER portions (green in **Fig. 2A**, Resting, upper panel and inset) within LAMP1-positive ELs in resting T cells (red circles in **Fig. 2A**, Resting, upper panel and inset). This testifies that basal ER-phagy operates in resting T cells. At day 1 p.a., the CANX-positive signal within the LAMP1-positive compartment was significantly reduced (from 2.0 ± 0.3 · 10^5^ arbitrary units (a.u.) in resting cells to 0.5 ± 0.1 · 10^5^ a.u., **Figs. 2A-B**, Resting *vs.* 1 Day).

**Figure 2.**
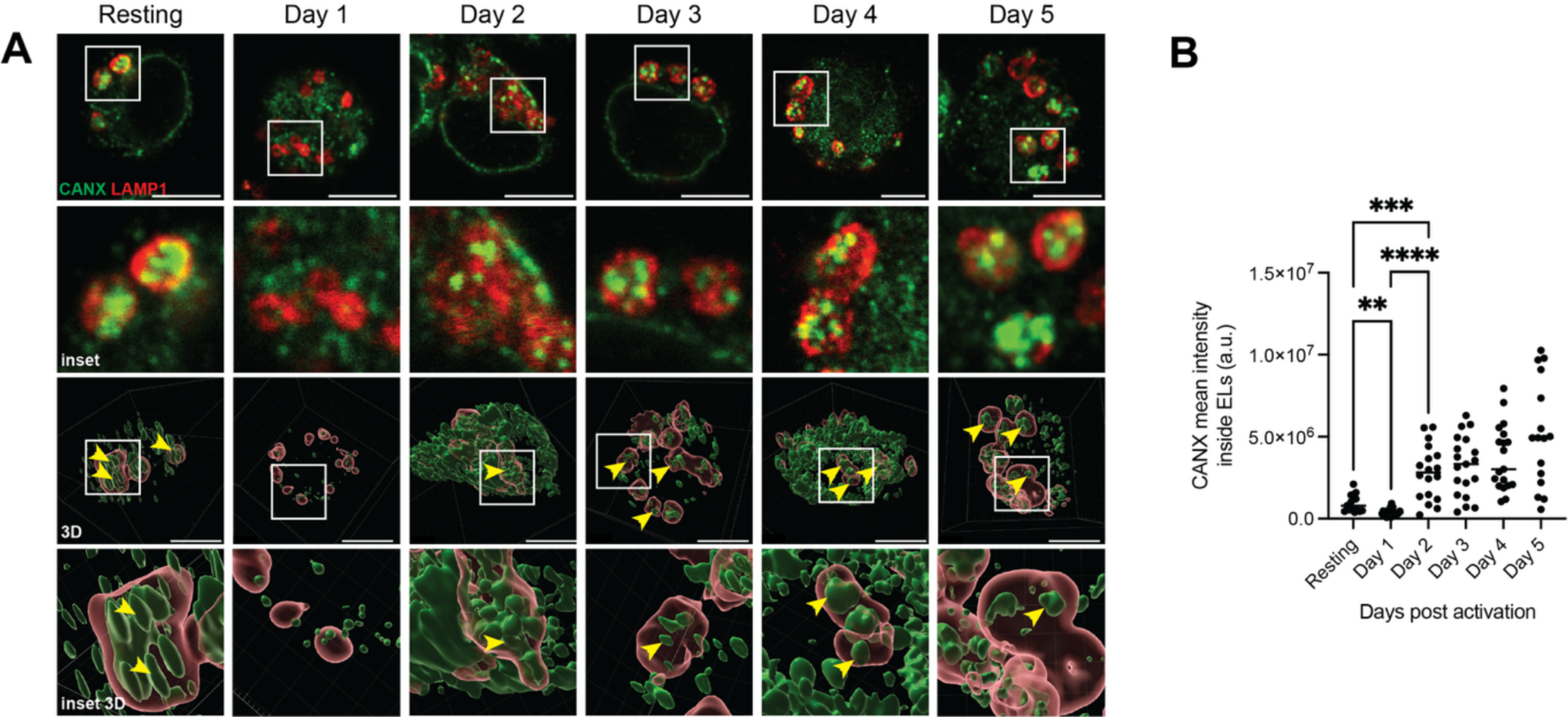
Delivery of CANX-positive ER portions to ELs in resting and activated CD4+ memory T cells as seen by LSCM. **A**) CANX accumulation inside ELs as monitored with LSCM in cells treated with BafA1 for 12 h before fixation. Top row: selected plane from Z-stack. Second row: insets. Third row: 3D reconstruction. Bottom row: insets of 3D reconstructions. Scale bars 5 µm. Insets 5×5 µm^2^. **B**) Quantification of average CANX intensity inside ELs (n= 16, 20, 18, 19, 19, 16 cells, respectively, from three separate experiments). One-way ANOVA and Dunnett’s multiple comparisons test, **P<0.01, ***P<0.001, ****P<0.0001.

Since the shutdown of lysosomal delivery of ER portions (**Fig. 2B**) occurred concomitantly with the initiation of cells’ enlargement (**Fig. 1B**) and with the transcriptional induction of ER-resident chaperones (**Figs. 1F, I, L**) but preceded the induction of ER chaperones’ proteins (**Figs. 1G-H, J-K, M-N**), we postulate that the inhibition of the constitutive ER turnover initiates the ER expansion, which will be eventually boosted by activation of the post-transcription phase (i.e., the production of ER-resident proteins) of the UPR. If this is correct, we anticipate that resting T cells with inactive ER-phagy are bigger or have larger ER content than resting T cells, where the ER-phagy program is normally executed. Lysosomal delivery of ER portions is eventually resumed from day 2 p.a., where an average value of 2.2 ± 0.5 · 10^5^ a.u. is measured (**Figs. 2A, 2B**). CANX intensity within ELs from day 2 p.a. up to day 5 p.a. tendentially increases, showing a possible role of ER-phagy in limiting the cellular enlargement resulting from the UPR activation, and in reducing the cell size during the contraction phase.

### Ultrastructural analyses of lysosomal delivery of ER portions in resting and in activated CD4+ memory T cells

Next, we performed Room Temperature Immuno-Electron Microscopy (RT-IEM) to obtain higher-resolution and temporal insight into the lysosomal delivery of the ER.

To achieve this, we developed a pre-embedding immunolabeling protocol for suspended cells, consisting of cells immobilization on glass coverslips by cytospin, sequential immunolabeling with anti-CANX and nanogold-conjugated secondary antibodies, gold enhancement, and flat embedding combined with serial-section (ss)TEM of 10 slices, 130-150nm each (see Methods for details). Using 3D ssRT-IEM, we evaluated the total CANX-positive ER portions delivered within ELs in all slices (**Fig. 3A** and Insets). CANX-associated gold particles were detected on ER membrane profiles in the cytoplasm (blue arrowheads, **Fig. 3A**, insets), or inside ELs (red arrowheads, **Fig. 3A**, insets). Some of the gold-labeled ER portions within the ELs are enclosed within larger vesicles (yellow arrowheads, **Fig. 3A**, insets), which would indicate that the ER portions have been captured by *micro*- or *macro*-ER-phagy programs (49). In resting cells, we counted, on average, 70.0 ± 16.2 CANX-associated gold particles per cell within all ELs. In agreement with the immunofluorescence 3D reconstructions (**Fig. 2A**), the CANX-associated gold particles per cell within all ELs were reduced to 20.4 ± 4.6 at day 1 p.a., to increase again to 142.0 ± 29.1 at day 2 p.a.

**Figure 3.**
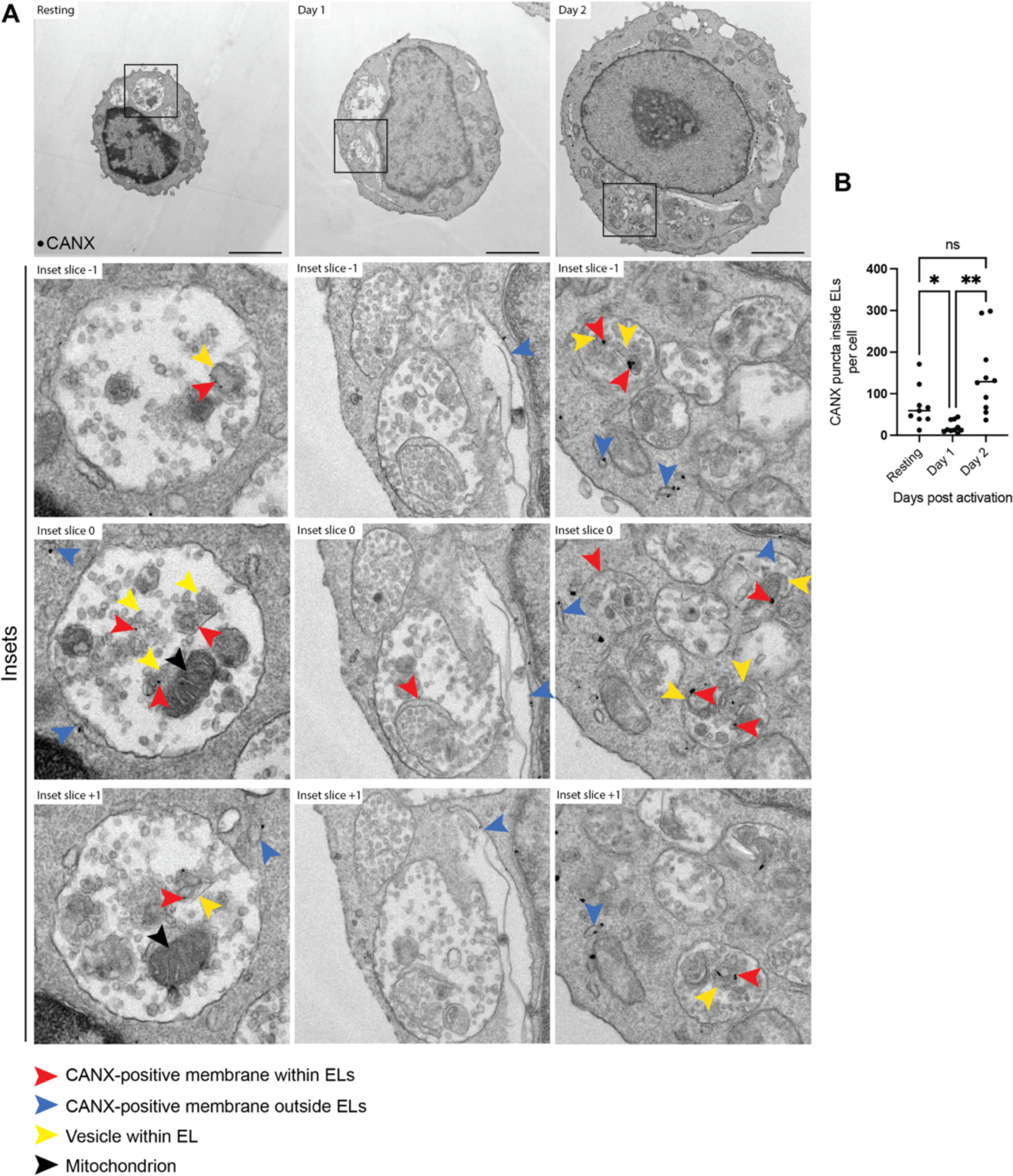
Delivery of CANX-positive ER portions to ELs in resting and activated CD4+ memory T cells as seen by ssRT-IEM. **A**) Top row: ssRT-IEM for CANX shows accumulation of ER portions within EL in resting cells treated with BafA1 for 12 h before fixation (Resting). Same for cells fixed 1 or 2 days p.a. (Day 1 and Day 2, respectively). Representative slice from reconstruction of 10 consecutive slices per cell. Bottom rows: Insets of 3 consecutive slices for each condition are shown. Scale bars 2 µm. Insets 2×2 μm^2^. **B**) Quantification of the number of CANX puncta inside ELs, subtracted for background (n=10 representative T cells). One-way ANOVA and Dunnett’s multiple comparisons test, ^ns^P>0.05, *P<0.05, **P<0.01.

### Lysosomal delivery of mitochondria portions in resting and in activated CD4+ memory T cells

Incidentally, the analyses of the content of the degradative compartments in resting T cells revealed the presence of mitochondrial portions (black arrowheads in **Fig. 3A**, Resting, insets) This prompted us to also explore the mitochondrial delivery within the LAMP1-positive ELs in T cells. To this end, we first performed immunofluorescence for the OMM marker TOMM20. As previously observed for the ER (**Fig. 2A**, Resting), analyses of resting T cells revealed the delivery of TOMM20-positive mitochondrial portions (blue in **Fig. 4A**, Resting first row and Insets in the second row) within LAMP1-positive ELs (red circles in **Fig. 4A** and Insets, **Fig. 4B**). At day 1 post-activation, the mitochondrial-associated immunofluorescence was virtually not detectable in the LAMP1-positive ELs (**Fig. 4A**, Day 1, and **Fig. 4B**). At later days p.a., the TOMM20-associated immunoreactivity was weakly detected again within the LAMP1-positive compartments, with a signal that appeared smaller, at the limit of detection for the confocal microscope, particularly with respect to the axial resolution and microscope’s optical sectioning capabilities (**Fig. 4A**, Days 2-5, and **Fig. 4B**).

**Figure 4.**
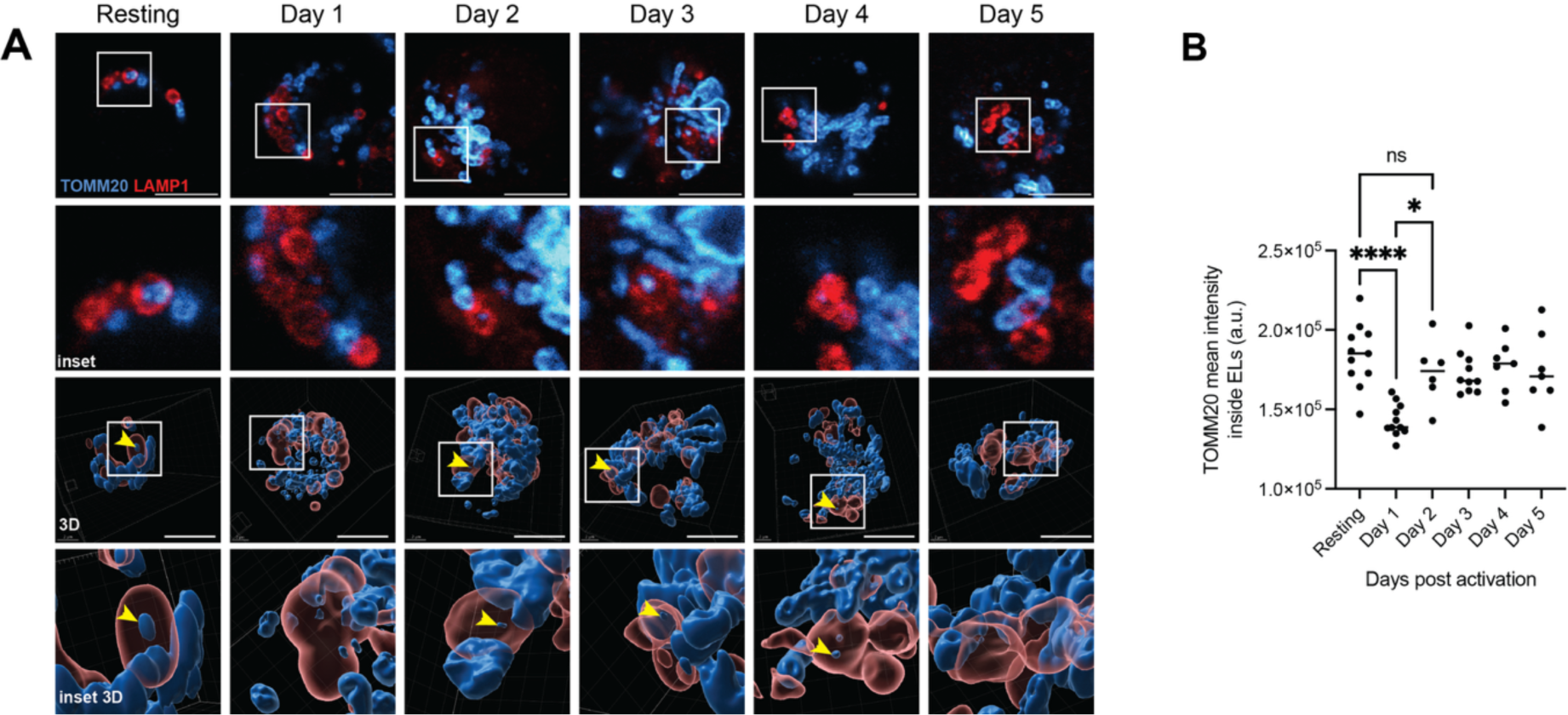
Delivery of TOMM20-positive mitochondrial portions to ELs in resting and activated CD4+ memory T cells as seen by LSCM. **A**) TOMM20 accumulation inside ELs as monitored by LSCM in cells treated with BafA1 for 12 h before fixation. Top row: selected plane from Z-stack. Second row: insets. Third row: 3D reconstruction. Bottom row: insets of 3D reconstructions. Scale bars 5 µm. Insets 5×5 µm^2^. **B**) Quantification of average TOMM20 intensity inside ELs (n=11, 11, 6, 10, 7, 7 cells respectively, from three separate experiments). One-way ANOVA and Dunnett’s multiple comparisons test, ^ns^P>0.05, *P<0.05, ****P<0.0001.

### Ultrastructural analyses of lysosomal delivery of mitochondrial portions in resting and in activated CD4+ memory T cells

To better investigate the TOMM20-positive signal accumulating within LAMP1-positive ELs in resting *vs.* activated T cells that we detected in the LSCM analyses, we also examined the intracellular distribution of the TOMM20-associated immunoreactivity by ssRT-IEM in BafA1-treated resting T cells and at day 1-2 p.a. (**Fig. 5A**). Analyses of the micrographs revealed that in resting cells gold-labeled TOMM20 was distributed at the limiting membrane of mitochondria in the cell cytoplasm (blue arrowheads, **Fig. 5A**, insets), characterized by an average volume of 6.8 ± 1.5 µm^3^ and of smaller mitochondrial portions with an average volume of 1.43 ± 0.41 µm^3^ contained within ELs (black arrowheads, **Fig. 5A**, Resting, insets). Notably, the mitochondrial portions accumulating within the inactive ELs of resting cells maintain the typical morphology characterized by an OMM and an IMM shaped in cristae (**Fig. 3A** and **Fig. 5A**, Resting, black arrowheads in the Insets). The micrographs of T cells 1 days p.a. confirmed the substantial reduction of the TOMM20-associated signal within the ELs as testified by the LSCM analyses (**Fig. 4A-B**). The micrographs of T cells 2 days p.a. also confirmed the LSCM analyses hinting at the lysosomal delivery of smaller TOMM20-positive structures, by revealing the presence of gold particles at the surface of single-membrane of mitochondria-derived vesicles lacking the conventional mitochondrial ultrastructure (green arrowheads, **Fig. 5A**, Day 2, insets) characterized by an average volume of 0.35 ± 0.11 µm^3^ (**Fig. 5C**).

**Figure 5.**
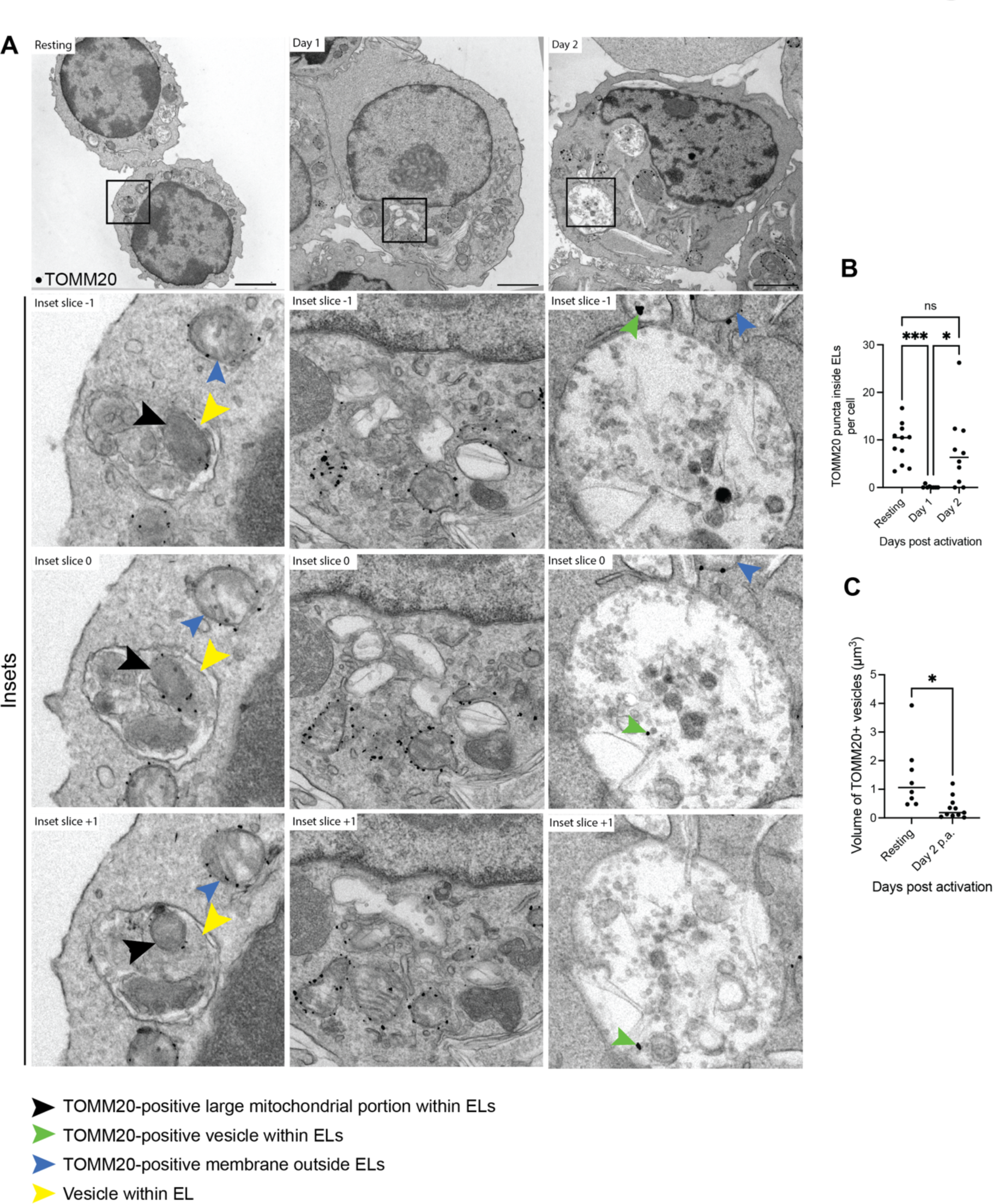
Delivery of TOMM20-positive mitochondrial portions to ELs in resting and activated CD4+ memory T cells as seen by ssRT-IEM. **A**) Top row: ssRT-IEM for TOMM20 shows accumulation of mitochondrial portions within EL in resting cells treated with BafA1 for 12 h before fixation (Resting). Same for cells fixed 1 or 2 days p.a. (Day 1 and Day 2, respectively). Representative slice from reconstruction of 10 consecutive slices per cell. Bottom rows: 3 consecutive slices for each condition are shown. Scale bars 2 µm. Insets 2×2 μm^2^. **B**) Quantification of the number of TOMM20 puncta inside ELs, subtracted for background (n=11, 10, 10 representative T cells respectively). One-way ANOVA and Dunnett’s multiple comparisons test, ^ns^P>0.05, *P<0.05, ***P<0.001. **C**) Quantification of the average volume of TOMM20+ vesicles, n=8. Mean line is shown. Unpaired two-tailed t-test, *P<0.05.

Collectively, LSCM and ssRT-IEM data show that human resting CD4+ T cells are characterized by a constitutive delivery of ER and mitochondrial material to the ELs, which is transiently shut down upon T cell activation to subsequently be resumed with distinct mechanism at least to regulate mitochondrial turnover.

## Discussion

In this study, we characterized the selective autophagy of the ER and mitochondria in human CD4+ memory T cells in the resting state and during activation. We report that resting human CD4+ memory T cells deliver CANX-positive ER portions to degradative LAMP-positive ELs for clearance. T cell activation is characterized by a transcriptional phase of the UPR showing a peak in ER stress genes mRNAs at day 1 p.a., and a somewhat delayed translational phase of the UPR characterized by a peak in ER stress proteins expression at 2-3 days p.a.. The translational UPR eventually triggers ER and cell expansion up to day 3 p.a. before induction of a contraction phase that gradually reduces the ER volume and the cell size. Interestingly, in the time lapse that separate the activation signal from the onset of the translational UPR phase testified by the increased levels of ER chaperone proteins, the T cells restrain the lysosomal delivery of ER and mitochondrial portions. We postulate that this contributes to the immediate increase of ER and mitochondrial content (upon inhibition of their constitutive lysosomal clearance) that sustain the first phases of T cells activation before the anabolic phase (the UPR for the ER and mtISR or mtUPR for mitochondria (13)) kicks in (**Fig. 6**, Day 0 to Day 1). Starting from Day 2, lysosomal delivery of ER and mitochondrial portions (in this phase OMM-derived vesicles) is resumed and slightly increases over time. Since the UPR and the mtISR are gradually vanishing, the overall balance of execution of the anabolic and catabolic programs eventually results in a contraction of cellular size (**Fig. 6**).

**Figure 6.**
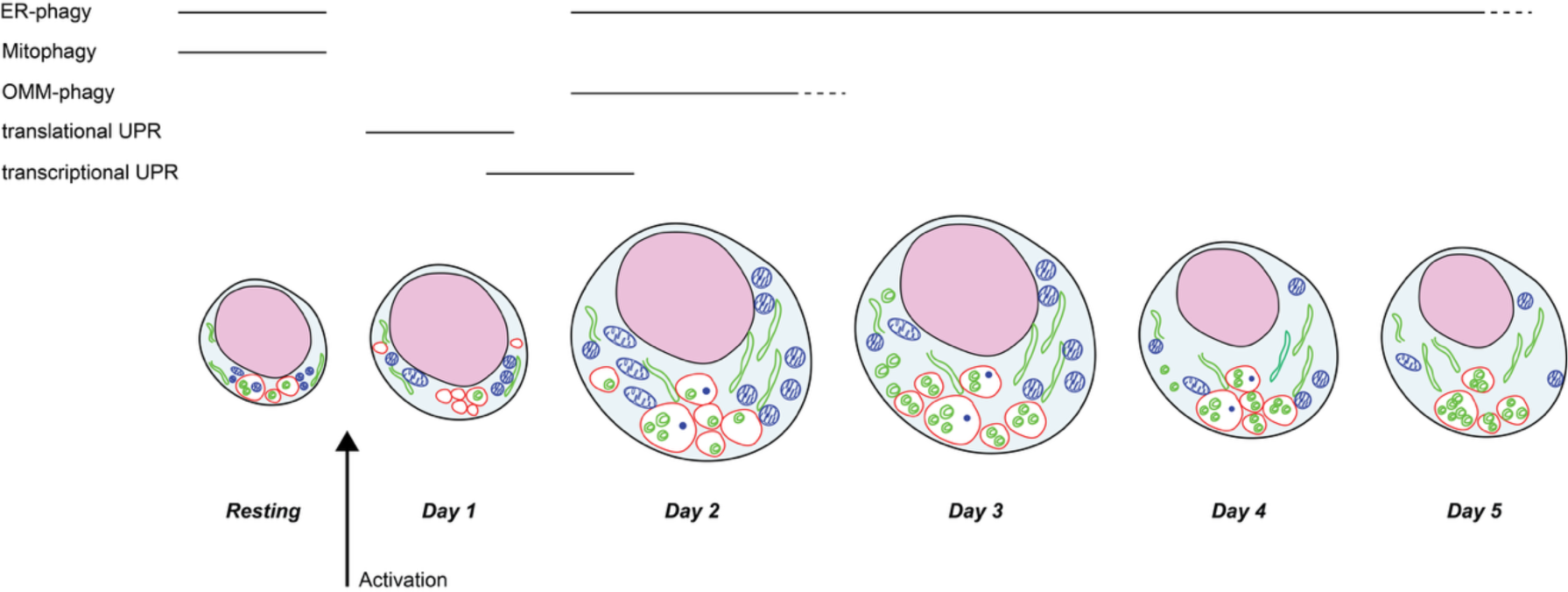
Graphical summary of the anabolic and catabolic processes in resting cells and after activation. CD4+ T cells remain in resting state for long times, during which they display active delivery of ER and mitochondrial portions to ELs for clearance. Immune challenge triggers cell activation, which consists of increase in cell and nucleus size, sustained cell growth and proliferation, activation of UPR, increase in ER to enhance the synthesis of protein, lipid and nucleic acids, and increase in total mitochondrial content. Upon activation, ER and mitochondrial delivery to ELs are transiently shutdown. From day 2 p.a., the ER is delivered again to ELs, as well as OMM-derived vesicles.

## Materials and Methods

### Reagents and Tools table

| Reagent/resource | Reference or source | Identifier or catalog number |
| --- | --- | --- |
| **Antibodies** |  |  |
| CD4+ positive selection beads | Miltenyi Biotec | 130-097-048 |
| Anti-CD3 | From A. Lanzavecchia |  |
| Anti-CD28 | BD Pharmigen | Clone CD28.2; 555725 |
| Anti-CD25-PE-Cy5 | Beckman Coulter | Clone B1.49.9; IM2646 |
| Anti-CD4-PE-Texas Red | Biolegend | Clone S3.5; MHCD0417 |
| Anti-CD45RA-Qdot 655 | Biolegend | Clone MEM-56; Q10069 |
| Anti-CCR7-BV421 | Biolegend | Clone G043H7; 353208 |
| Rabbit anti-Calnexin | From A. Helenius |  |
| Rabbit anti-BiP | Cell Signaling | 3183 |
| Rabbit anti-HERP | From K. Kokame |  |
| Mouse anti-LAMP1 | DSHB (J.T. August) | Clone H4A3; 1D4B |
| Rabbit anti-TOMM20 | Abcam | Clone EPR-155891-39; Ab186734 |
| **Chemicals, enzymes and other reagents** |  |  |
| SPY650-DNA | Spirochrome |  |
| Bafilomycin A1 | Sigma Aldrich | B1793 |
| Abberior Mount | Abberior GmbH | MM-2009 |
| GenElute Mammalian Total RNA Miniprep Kit | Sigma | RTN70 |
| SuperScript II reverse transcriptase | Thermo Fisher | 18064014 |
| Power SYBR Green PCR Master Mix | Thermo Fisher | 4367659 |
| **Software** |  |  |
| IDEAS 6.4 | Cytek |  |
| Fiji/ImageJ | https://imagej.nih.gov/ij/index.html |  |
| LysoQuant plugin for ImageJ | Morone et al. 2020 |  |
| Velox 3.6.0 | Thermo Fisher |  |
| MAGE3D | https://wp.unil.ch/emf/software-mage3d/ |  |
| GraphPad 10 | https://www.graphpad.com |  |
| QuantStudio | QuantStudio™ Design & Analysis Software v1.5.5 |  |
| **Other** |  |  |
| ImageStream X Mark II | Cytek |  |
| Leica Stellaris 5 | Leica |  |
| Leica Stellaris 8 STED FALCON | Leica |  |
| Talos L120C | Thermo Fisher |  |
| QuantStudio™ 3 RT-PCR | Thermo Fisher |  |

### Cells purification and sorting

Blood from healthy donors was obtained from the Swiss Blood Donation Center of Basel and Lugano, with informed consent from the Swiss Red Cross and authorization number CE 3428 from the Comitato Etico Canton Ticino. CD4+ T cells were separated by gradient centrifugation (Ficoll-Paque Plus; GE Healthcare), followed by positive selection using magnetic microbeads (Miltenyi Biotec). After washing, cells were stained on ice for 30 min with the following fluorochrome-labeled mouse monoclonal antibodies: Naive and memory CD4+ T cells were sorted to >98% purity on a FACS S6 Sorter (BD) after exclusion of CD25bright cells. Naive T cells were sorted as CD4+CD45RA+CCR7+CD95−; the remaining CD4+ T cells were sorted as total memory cells.

### Cell culture

T cells were cultured in RPMI 1640 supplemented with 2 mM glutamine, 1% (vol/vol) nonessential amino acids, 1% (vol/vol) sodium pyruvate, penicillin (50 U/ml), streptomycin (50 μg/ml; all from Invitrogen), and 5% human serum (Swiss Red Cross). For activation experiments, T lymphocytes were stimulated in NUNC 96-well plates (ThermoFisher) using plate-bound anti-CD3 (1 μg ml^−1^, clone TR66, recombinant, made in-house) and anti-CD28 (1 μg ml^−1^) and medium was supplemented with IL-2 (500 IU/ml). BafA1 (Sigma) was used at 100 mM for 12 h or 50 mM for 6 h.

### Antibodies

Antibodies used in this study are anti-CD3 (recombinant, kind gift of A. Lanzavecchia), anti-CD28 (clone CD28.2; 555725; BD Pharmigen), anti-CD25-PE-Cy5 (clone B1.49.9; IM2646; Beckman Coulter), anti-CD4-PE-Texas Red (clone S3.5; MHCD0417; Biolegend), anti-CD45RA-Qdot 655 (clone MEM-56; Q10069; Biolegend), and anti-CCR7-BV421 (clone G043H7; 353208; Biolegend), anti-LAMP1 (Hybridoma Bank, 1D4B deposited by J.T. August), anti-BiP (clone 3183; Cell Signaling), anti-HERP (kind gift of K. Kokame), anti-CANX (kind gift A. Helenius). Alexa-conjugated secondary antibodies (Alexa Fluor 488 and Alexa Fluor 568) were purchased from Thermo Fisher Scientific.

### Imaging Flow Cytometry (IFC)

Cells were fixed with 4% PFA diluted in PBS for 20 minutes at RT in suspension. Cells were permeabilized for 15 min with permeabilization solution (PS) (0.05% saponin, 10% goat serum, 10mM HEPES, 15mM glycine). Primary antibodies were diluted 1:100 in PS for 90 min, washed for 15 min in PBS, and then incubated with Alexa Fluor-conjugated secondary antibodies diluted 1:300 in PS for 45 min. SPY650-DNA nuclear staining (Spirochrome) was added to last wash at 1:5000 dilution. Images were acquired at ImageStream X Mark II cytometer (Cytek) using an Olympus 60x/0.9NA objective and 488 nm, 561 nm and 638 nm lasers. Flow was set to low speed for maximum resolution and EDF (Extended Depth of Field) was activated. Analysis was performed with IDEAS 6.4 software (Cytek).

### Laser Scanning Confocal Microscopy (LSCM)

Cells were fixed with 4% PFA diluted in PBS for 20 minutes at RT in suspension and then cytospinned at 1500rpm for 5 minutes on a 12 mm #1.5 thickness coverslip (Cytospin 4, Thermo Fisher). Cells were permeabilized for 15 min with permeabilization solution (PS) (0.05% saponin, 10% goat serum, 10 mM HEPES, 15 mM glycine). Primary antibodies were diluted 1:100 in PS for 90 min, washed for 15 min in PS, and then incubated with Alexa Fluor-conjugated secondary antibodies diluted 1:300 in PS for 45 min. Cells were rinsed with PS and water and mounted with Abberior Mount (Abberior) or Vectashield (Vector Labs) with or without 4’,6-diamidino-2-phenylindole (DAPI). Images were acquired on a Leica Stellaris 8 STED FALCON or Leica Stellaris 5 (Leica), with a Plan Apo 63x/1.4NA objective, excitations 405 nm, 499 nm and 579 nm and emission ranges 430-494 nm, 504-573 nm and 585-740 nm respectively. Pinhole was set to 0.8 AU (estimated optical sectioning of about 0.8 μm). Images were acquired with a pixel dwell time of 2.8 μs, at a voxel size of 40 nm x 230 nm. Deconvolution was performed with LAS X (Leica). Images were quantified with ImageJ and LysoQuant (Morone et al. 2020) or Imaris 10.2 (Andor/Bitplane).

### Serial section Room-Temperature Immunogold Electron Microscopy (ssRT-IEM)

Cells were fixed in 4% PFA EM grade and 0.2 M HEPES buffer for 1h at RT or in Periodate-lysine-paraformaldehyde (PLP) for 2 h at RT. Cells were resuspended in PBS and cytospinned as above on glass coverslips. After three washes in PBS, cells were incubated 10 min with 50 mM glycine and blocked 1 h in blocking buffer (0.2% bovine serum albumin, 5% goat serum, 50 mM NH4Cl, 0.1% saponin, 20 mM PO4 buffer, 150 mM NaCl). Staining with primary antibodies and nanogold-labeled secondary antibodies (Nanoprobes) were performed in a blocking buffer at RT. Cells were fixed for 30 min in 1% GA and nanogold was enlarged with gold enhancement solution (Nanoprobes) according to the manufacturer’s instructions. After several washes in cacodylate buffer, cells were post-fixed in 1% osmium tetroxide (OsO4), 1.5% potassium ferricyanide (K4[Fe(CN)6]) in 0.1 M Na-cacodylate buffer for 1 h on ice, washed with distilled water (dH2O) and *en bloc* stained with 0.5% uranyl acetate in dH2O overnight at 4 °C in the dark. Samples were rinsed in dH2O, dehydrated with increasing concentrations of ethanol, embedded in Epon resin and cured in an oven at 60 °C for 48 h. Ultrathin sections (70–90 nm, or 130-150 nm) were collected on formvar carbon-coated slot grids using an ultramicrotome (UC7, Leica microsystem, Vienna, Austria), stained with uranyl acetate and Sato’s lead solutions and observed in a Transmission Electron Microscope Talos L120C (FEI, Thermo Fisher Scientific) operating at 120 kV. Images were acquired with a Ceta CCD camera (FEI, Thermo Fisher Scientific) using Velox 3.6.0 (FEI, Thermo Fisher Scientific). Serial sections were reconstructed with MAGE3D program (University of Lausanne) and analyzed with ImageJ. To subtract gold background from puncta quantification, IEM gold spots were counted in ELs and in Nucleus for each cell section, and the volume of acquired ELs and Nucleus were measured (area times slice thickness). Total puncta density in the nucleus (staining background) was then subtracted from EL puncta density, and this density was multiplied by EL volume to obtain final, background-subtracted number of puncta in the ELs.

### RNA extraction, polymerase chain reaction with reverse transcription (RT–PCR)

The extraction of RNA was performed with the GenElute Mammalian Total RNA Miniprep Kit (Sigma) according to the manufacturer’s instructions. One microgram of RNA was used for cDNA synthesis with dNTPs (Kapa Biosystems), oligo(dT) and the SuperScript II reverse transcriptase (Thermo Fisher Scientific) according to the instructions of the manufacturer. For each qRT-PCR reaction, 10 μl of Power SYBR Green PCR Master Mix (Bimake), 0.4 μl of reference ROX dye and 7.6 μl of milliQ sterile water were added to 1 μl cDNA together with 1 μl of 10 μM forward and reverse primer mix human (h)BiP for 5’-GAG TTC TTC AAT GGC AAG GA-3’, rev 5’-CCA GTC AGA TCA AAT GTA CCC-3’, hHERPUD1 for 5’-ATG GAG TCC GAG ACC GAA C-3’, rev 5’-TTG GTG ATC CAA CAA CAG CTT-3’ hCANX for 5’-CCA AGC ATC ATG CCA TCT CT-3’, rev 5’-TTG GTC TTT CAT CCC AAT CC-3’, hSDHA for 5’-ACA GAG CCT CAA GTT TGC AAA G-3’, rev 5’-AAA TAG CTG GTA TCA TAT CGC AGA G-3’) for the transcript of interest in 96-well reaction plate (MicroAmp Fast Optical 96-Well Reaction Plate with Barcode (0.1 ml), Applied Biosystems). The plate was vortexed and centrifuged. Samples were loaded as triplicates. Quantitative real-time PCR was performed using QuantStudio™ 3 Real-Time PCR System. The housekeeping gene actin was used as reference. Data were analyzed using the QuantStudio™ Design & Analysis Software v1.5.5.

### Cell lysis and Western blot

After the respective treatments, 1 million cells per condition were washed with ice-cold PBS containing 20 mM N-ethylmaleimide (NEM) then lysed with RIPA buffer (1% Triton X-100, 0.1% SDS, 0.5% sodium deoxycholate in HBS, pH 7.4) supplemented with protease inhibitors (1 mM phenylmethylsulfonyl fluoride (PMSF), 16.5 mM Chymostatin, 23.4 mM Leupeptin, 16.6 mM Antipain, 14.6 mM Pepstatin). PNS were collected after centrifugation at 10,600 g, 4 °C for 10 min, and reduced by adding 100 mM dithiothreitol (DTT; F. Hoffmann-La Roche AG, Basel, Switzerland) and heating at 95 °C for 5 min, then subjected to SDS–PAGE. Proteins were transferred to PVDF membranes using the Trans-Blot Turbo Transfer System (Bio-Rad). For Western blotting (WB), proteins were transferred from polyacrylamide gels to polyvinylidene fluoride (PVDF) membranes using the TransBlot Turbo device (Bio-Rad Laboratories AG, Cressier, Fribourg, Switzerland). The PVDF membranes were blocked for 10 min with 10% milk (BioRad, w/v) in Tris-buffered saline and 0.1% Tween 20 (TBS-T), briefly rinsed in TBS-T, and incubated overnight at 4 °C with primary antibodies under agitation. After washing out the primary antibodies with TBS-T, membranes were incubated with HRP-conjugated secondary antibodies or protein A for 45 min at RT with shaking. Protein bands were detected using the Fusion FX7 chemiluminescence detection system (Vilber, Collégien (Marne-la-Vallée), France) and the WesternBright™ Quantum system (Advansta Inc., San Jose, CA, USA), according to the manufacturer’s instructions. Protein band intensities were quantified with ImageJ software (U.S. National Institutes of Health, Bethesda, MD, USA).

### Statistical analysis

Plots and statistical analyses were performed using GraphPad Prism 10 (GraphPad Software Inc.). In this study, one-way ANOVA with Dunnett’s multiple comparisons test and unpaired two-tailed t-test were used to assess statistical significance. An adjusted P-value < 0.05 was considered as statistically significant (* P<0.05, **, P<0.01, *** P<0.001, *** P<0.0001).

## Acknowledgments

We would like to thank the members of Molinari’s lab and David Jarrossay for discussions and critical reading of the manuscript. We thank the members of S. Monticelli’s lab for cells and materials received, A. Lanzavecchia, K. Kokame and A. Helenius for materials received.

## Funding

Swiss National Science Foundation grants 310030_214903 and 320030-227541 (MM). Alpha1-Foundation Application ID: 1188512. IRB’s pilot promising projects (PPP) funding.

## Notes

### Competing Interest Statement

The authors have declared no competing interest.

## References

1. Lam N, Lee Y, Farber DL. A guide to adaptive immune memory. Nat Rev Immunol (2024) doi: 10.1038/s41577-024-01040-6

2. Lam N, Angel JC, Buchholz BA, Lee Y, Weisberg SP, Brown BH, Davis-Porada J, Caron DP, Jensen IJ, Szabo PA, et al. Asynchronous aging and turnover of human circulating and tissue-resident memory T cells across sites. Immunity (2025) 0: doi: 10.1016/j.immuni.2025.07.001

3. Hwang J-R, Byeon Y, Kim D, Park S-G. Recent insights of T cell receptor-mediated signaling pathways for T cell activation and development. Exp Mol Med (2020) 52:750–761. doi: 10.1038/s12276-020-0435-8

4. Künzli M, Masopust D. CD4+ T cell memory. Nat Immunol (2023) 24:903–914. doi: 10.1038/s41590-023-01510-4

5. Schwarz DS, Blower MD. The endoplasmic reticulum: structure, function and response to cellular signaling. Cell Mol Life Sci (2016) 73:79–94. doi: 10.1007/s00018-015-2052-6

6. Hetz C, Zhang K, Kaufman RJ. Mechanisms, regulation and functions of the unfolded protein response. Nat Rev Mol Cell Biol (2020) 21:421–438. doi: 10.1038/s41580-020-0250-z

7. Preissler S, Ron D. Early Events in the Endoplasmic Reticulum Unfolded Protein Response. Cold Spring Harb Perspect Biol (2019) 11:a033894. doi: 10.1101/cshperspect.a033894

8. Kemp K, Poe C. Stressed: The Unfolded Protein Response in T Cell Development, Activation, and Function. International Journal of Molecular Sciences (2019) 20:1792. doi: 10.3390/ijms20071792

9. Zhang W, Cao X. Unfolded protein responses in T cell immunity. Front Immunol (2025) 15:1515715. doi: 10.3389/fimmu.2024.1515715

10. Pengo N, Scolari M, Oliva L, Milan E, Mainoldi F, Raimondi A, Fagioli C, Merlini A, Mariani E, Pasqualetto E, et al. Plasma cells require autophagy for sustainable immunoglobulin production. Nat Immunol (2013) 14:298–305. doi: 10.1038/ni.2524

11. Cluxton D, Petrasca A, Moran B, Fletcher JM. Differential Regulation of Human Treg and Th17 Cells by Fatty Acid Synthesis and Glycolysis. Front Immunol (2019) 10: doi: 10.3389/fimmu.2019.00115

12. Franco A, Almanza G, Burns JC, Wheeler M, Zanetti M. Endoplasmic reticulum stress drives a regulatory phenotype in human T-cell clones. Cellular Immunology (2010) 266:1–6. doi: 10.1016/j.cellimm.2010.09.006

13. Suomalainen A, Nunnari J. Mitochondria at the crossroads of health and disease. Cell (2024) 187:2601–2627. doi: 10.1016/j.cell.2024.04.037

14. Buck MD, O’Sullivan D, Klein Geltink RI, Curtis JD, Chang C-H, Sanin DE, Qiu J, Kretz O, Braas D, van der Windt GJW, et al. Mitochondrial Dynamics Controls T Cell Fate through Metabolic Programming. Cell (2016) 166:63–76. doi: 10.1016/j.cell.2016.05.035

15. Chapman NM, Boothby MR, Chi H. Metabolic coordination of T cell quiescence and activation. Nat Rev Immunol (2020) 20:55–70. doi: 10.1038/s41577-019-0203-y

16. Norton EG, Chapman NM, Chi H. Mitochondria and lysosomes in T cell immunometabolism. Trends in Immunology (2025) 46:635–651. doi: 10.1016/j.it.2025.07.014

17. Śniegocka M, Liccardo F, Fazi F, Masciarelli S. Understanding ER homeostasis and the UPR to enhance treatment efficacy of acute myeloid leukemia. Drug Resist Updat (2022) 64:100853. doi: 10.1016/j.drup.2022.100853

18. Lin Y-F, Haynes CM. Metabolism and the UPR(mt). Mol Cell (2016) 61:677–682. doi: 10.1016/j.molcel.2016.02.004

19. Oakes SA, Papa FR. The role of endoplasmic reticulum stress in human pathology. Annu Rev Pathol (2015) 10:173–194. doi: 10.1146/annurev-pathol-012513-104649

20. Gergely Jr. P, Grossman C, Niland B, Puskas F, Neupane H, Allam F, Banki K, Phillips PE, Perl A. Mitochondrial hyperpolarization and ATP depletion in patients with systemic lupus erythematosus. Arthritis & Rheumatism (2002) 46:175–190. doi: 10.1002/1529-0131(200201)46:1%3C175::AID-ART10015%3E3.0.CO;2-H

21. Reggiori F, Molinari M. ER-phagy: mechanisms, regulation, and diseases connected to the lysosomal clearance of the endoplasmic reticulum. Physiological Reviews (2022) 102:1393–1448. doi: 10.1152/physrev.00038.2021

22. Hoyer MJ, Capitanio C, Smith IR, Paoli JC, Bieber A, Jiang Y, Paulo JA, Gonzalez-Lozano MA, Baumeister W, Wilfling F, et al. Combinatorial selective ER-phagy remodels the ER during neurogenesis. Nat Cell Biol (2024) 26:378–392. doi: 10.1038/s41556-024-01356-4

23. Vats S, Galli T. Introducing secretory reticulophagy/ER-phagy (SERP), a VAMP7-dependent pathway involved in neurite growth. Autophagy (2021) 17:1037–1039. doi: 10.1080/15548627.2021.1883886

24. Yao X, Wu Y, Xiao T, Zhao C, Gao F, Liu S, Tao Z, Jiang Y, Chen S, Ye J, et al. T-cell-specific Sel1L deletion exacerbates EAE by promoting Th1/Th17-cell differentiation. Molecular Immunology (2022) 149:13–26. doi: 10.1016/j.molimm.2022.06.001

25. Sinclair LV, Youdale T, Spinelli L, Gakovic M, Langlands AJ, Pathak S, Howden AJM, Ganley IG, Cantrell DA. Autophagy repression by antigen and cytokines shapes mitochondrial, migration and effector machinery in CD8 T cells. Nat Immunol (2025) 26:429–443. doi: 10.1038/s41590-025-02090-1

26. Uoselis L, Nguyen TN, Lazarou M. Mitochondrial degradation: Mitophagy and beyond. Molecular Cell (2023) 83:3404–3420. doi: 10.1016/j.molcel.2023.08.021

27. Denk D, Petrocelli V, Conche C, Drachsler M, Ziegler PK, Braun A, Kress A, Nicolas AM, Mohs K, Becker C, et al. Expansion of T memory stem cells with superior anti-tumor immunity by Urolithin A-induced mitophagy. Immunity (2022) 55:2059–2073.e8. doi: 10.1016/j.immuni.2022.09.014

28. Bektas A, Schurman SH, Gonzalez-Freire M, Dunn CA, Singh AK, Macian F, Cuervo AM, Sen R, Ferrucci L. Age-associated changes in human CD4+ T cells point to mitochondrial dysfunction consequent to impaired autophagy. Aging (2019) 11:9234–9263. doi: 10.18632/aging.102438

29. Liu X, Yu J, Xu L, Umphred-Wilson K, Peng F, Ding Y, Barton BM, Lv X, Zhao MY, Sun S, et al. Notch-induced endoplasmic reticulum-associated degradation governs mouse thymocyte β−selection. eLife (2021) 10:e69975. doi: 10.7554/eLife.69975

30. Richter FC, Saliutina M, Hegazy AN, Bergthaler A. Take my breath away— mitochondrial dysfunction drives CD8+ T cell exhaustion. Genes Immun (2024) 25:4–6. doi: 10.1038/s41435-023-00233-8

31. Zhang R, Gao F, Li J, Jin J, Chen K, Chaudhuri S, Liao Z, Xiao T, Xu Y, Wen H, et al. USP30 inhibition augments mitophagy to prevent T cell exhaustion. Science Advances (2025) 11:eadv6902. doi: 10.1126/sciadv.adv6902

32. Franco F, Bevilacqua A, Wu R-M, Kao K-C, Lin C-P, Rousseau L, Peng F-T, Chuang Y-M, Peng J-J, Park J, et al. Regulatory circuits of mitophagy restrict distinct modes of cell death during memory CD8+ T cell formation. Sci Immunol (2023) 8:eadf7579. doi: 10.1126/sciimmunol.adf7579

33. Luo L, Lechuga-Vieco AV, Sattentau C, Borsa M, Simon AK. Dysfunctional mitochondria in ageing T cells: a perspective on mitochondrial quality control mechanisms. EMBO reports (2025) 26:4402–4418. doi: 10.1038/s44319-025-00536-z

34. Wang N, Yuan J, Karim MR, Zhong P, Sun Y-P, Zhang H-Y, Wang Y-F. Effects of Mitophagy on Regulatory T Cell Function in Patients With Myasthenia Gravis. Front Neurol (2020) 11: doi: 10.3389/fneur.2020.00238

35. Borsa M, Lechuga-Vieco AV, Kayvanjoo AH, Yazicioglu Y, Compeer EB, Richter FC, Bui H, Dustin ML, Katajisto P, Simon AK. Inheritance of old mitochondria controls early CD8 ^+^ T cell fate commitment and is regulated by autophagy. (2024) doi: 10.1101/2024.01.29.577412

36. Duthoo E, Vral A, Baeyens A. An updated view into the cell cycle kinetics of human T lymphocytes and the impact of irradiation. Sci Rep (2022) 12:7687. doi: 10.1038/s41598-022-11364-9

37. Saravia J, Chapman NM, Chi H. Helper T cell differentiation. Cell Mol Immunol (2019) 16:634–643. doi: 10.1038/s41423-019-0220-6

38. Cornish GH, Sinclair LV, Cantrell DA. Differential regulation of T-cell growth by IL-2 and IL-15. Blood (2006) 108:600–608. doi: 10.1182/blood-2005-12-4827

39. Waugh RE, Lomakina E, Amitrano A, Kim M. Activation effects on the physical characteristics of T lymphocytes. Front Bioeng Biotechnol (2023) 11: doi: 10.3389/fbioe.2023.1175570

40. Low JS, Vaqueirinho D, Mele F, Foglierini M, Jerak J, Perotti M, Jarrossay D, Jovic S, Perez L, Cacciatore R, et al. Clonal analysis of immunodominance and cross-reactivity of the CD4 T cell response to SARS-CoV-2. Science (2021) 372:1336– 1341. doi: 10.1126/science.abg8985

41. Forrester A, De Leonibus C, Grumati P, Fasana E, Piemontese M, Staiano L, Fregno I, Raimondi A, Marazza A, Bruno G, et al. A selective ER-phagy exerts procollagen quality control via a Calnexin-FAM134B complex. The EMBO Journal (2019) 38:e99847. doi: 10.15252/embj.201899847

42. Wang M, Wey S, Zhang Y, Ye R, Lee AS. Role of the Unfolded Protein Response Regulator GRP78/BiP in Development, Cancer, and Neurological Disorders. Antioxid Redox Signal (2009) 11:2307–2316. doi: 10.1089/ars.2009.2485

43. Pfaffenbach KT, Lee AS. The critical role of GRP78 in physiologic and pathologic stress. Current Opinion in Cell Biology (2011) 23:150–156. doi: 10.1016/j.ceb.2010.09.007

44. Kokame K, Kato H, Miyata T. Identification of ERSE-II, a New cis-Acting Element Responsible for the ATF6-dependent Mammalian Unfolded Protein Response. Journal of Biological Chemistry (2001) 276:9199–9205. doi: 10.1074/jbc.m010486200

45. Geigges M, Gubser PM, Unterstab G, Lecoultre Y, Paro R, Hess C. Reference Genes for Expression Studies in Human CD8+ Naïve and Effector Memory T Cells under Resting and Activating Conditions. Sci Rep (2020) 10:9411. doi: 10.1038/s41598-020-66367-1

46. Klionsky DJ, Elazar Z, Seglen PO, Rubinsztein DC. Does bafilomycin A _1_ block the fusion of autophagosomes with lysosomes? Autophagy (2008) 4:849–850. doi: 10.4161/auto.6845

47. Fregno I, Fasana E, Bergmann TJ, Raimondi A, Loi M, Soldà T, Galli C, D’Antuono R, Morone D, Danieli A, et al. ER-to-lysosome-associated degradation of proteasome-resistant ATZ polymers occurs via receptor-mediated vesicular transport. The EMBO journal (2018)e99259. doi: 10.15252/embj.201899259

48. Morone D, Marazza A, Bergmann TJ, Molinari M. Deep learning approach for quantification of organelles and misfolded polypeptides delivery within degradative compartments. MBoC (2020)mbc.E20-04-0269. doi: 10.1091/mbc.E20-04-0269

49. Loi M, Raimondi A, Morone D, Molinari M. ESCRT-III-driven piecemeal micro-ER-phagy remodels the ER during recovery from ER stress. Nature Communications (2019) 10: doi: 10.1038/s41467-019-12991-z

